# *Present*: flexible neuroscience- and technology-driven frameworks for the study of classroom engagement

**DOI:** 10.1101/2023.06.23.544732

**Authors:** Conor Lee Shatto, John Thorp, Josh Sterling Friedman, Isabella Rosario, Zhuowei Gu, Junsheng Shi, Megan Zhuo, Adam Brown, Xiaofu He, Alfredo Spagna

## Abstract

Classroom engagement’s impact on academic success is crucial. However, the contributions of affective, cognitive, and behavioral components of engagement remain uncertain. We conducted two studies using non-invasive, research-based approaches to clarify these contributions. Study 1 employed portable EEG headsets to measure cognitive engagement, in-class quizzes assessed content retention, and post-class subjective questionnaires indexed affective engagement by measuring feelings of learning and engagement. Content retention predicted subjective measures, while the EEG theta/beta ratio was negatively related to content retention but positively related to subjective measures. Study 2 featured embedded measures of content retention, confidence, engagement, background knowledge, and indexed behavioral engagement looking at nonverbal behavior quantified via video camera recordings. Confidence and engagement were significantly correlated with each other and with particular facial muscle, gaze direction, and head pose movements. We discuss how these approaches enable real-time studies of classroom engagement and can be integrated to develop neurofeedback interventions.

## Introduction

Traditionally, scores on midterms and final exams are assumed to be proxies for student skill attainment and knowledge retention. However, such post-hoc evaluations have been criticized for inducing performance-hindering anxiety (von der Embse et al., 2018) and failing to accurately predict academic readiness (Allensworth & Clark, 2020). Meanwhile, classroom engagement has long been considered among the most important predictors of successful learning and academic achievement (Alyahyan & Düştegör, 2020; Marks, 2000; Newmann, 1992). A detailed understanding of classroom engagement and its relationship with skills attainment and knowledge retention is therefore critical for pedagogical innovation and development of academic interventions within and outside the classroom.

Classroom engagement is a multidimensional and multi-temporal construct that encompasses affective, cognitive, and behavioral aspects, and spans time scales from seconds (micro-levels) to hours (macro-levels; D’Mello et al., 2017). Behavioral engagement comprises the overt actions and participation of students in classroom activities; as the most directly observable, it is the most widely studied of the three types (Suárez-Orozco et al., 2009) and has been shown to predict academic achievement in informal educational settings (Fredricks et al., 2004) and ecological classroom contexts (Lane & Harris, 2015). Constructs of behavioral engagement include on-task behaviors, such as paying attention or contributing to class discussion, general attendance, and participation in academic activities outside the classroom (Hospel et al., 2016). Affective engagement refers to students’ emotional reactions to learning activities (Connell & Wellborn, 1991; Skinner & Belmont, 1993). Broadly, it encompasses students’ interest, enjoyment, enthusiasm, boredom, and anxiety for learning. Motivational constructs, such as a sense of belonging with peers or within the larger school system are often categorized as affective and are related to academic success (Fredricks & McColskey, 2012; Khan et al., 2023; Wara et al., 2018). Lastly, cognitive engagement refers to metacognitive and self-regulatory strategies that students use to better comprehend instructional material (Flavell, 1979). Encompassing the psychological, covert processes of learning and cognitive strategies, such as level of persistence, effort exerted towards academic activities, and attention (Lam et al., 2014), cognitive engagement is highly predictable of successful learning (Greene et al., 2004). Furthermore, high cognitive engagement has been linked to enhanced motivation (Guthrie et al., 2004), self-regulation (Cleary & Zimmerman, 2012), and self-efficacy (Walker et al., 2006).

As noted above, a wealth of evidence supports the individual connection between each of the three components of classroom engagement and academic success. A comprehensive characterization of the synergistic contributions of these components in supporting academic success is still lacking, as most studies have examined only one component in isolation (e.g., facial expressions as a proxy of emotional engagement); (Grafsgaard et al., 2013). The simultaneous study of all components of classroom engagement requires fitting multiple factor models and collection methods into the same study protocol (Shernof et al., 2017), but methods to reliably measure engagement’s many components simultaneously have not yet been developed (Betts, 2012). Without a simultaneous characterization, it is difficult to identify the causes of naturally occurring engagement fluctuations, or develop effective real-time interventions that promote engagement in the classroom (Fredricks & Eccles, 2006).

The field of education research is increasingly focused on developing a holistic conceptualization of classroom engagement that integrates these components into a single framework. Recent results indicate that cognitive neuroscience methods may provide a backbone to build this new, multifaceted approach to studying classroom engagement (Davidesco, 2020; Davidesco et al., 2023). Neuroscience research has traditionally taken place in tightly controlled laboratory settings, featuring expensive and cumbersome measurement tools. At the same time, neuroscience is uniquely suited at the conceptual level to develop evidence-based frameworks for the integrative study of classroom engagement by leveraging its rich models of attention, memory, emotional regulation, and executive functions that support and affect learning. Capitalizing on recent technological advances in order to bridge this divide between methodological constraints and conceptual opportunities, the *neuroscience of education* approach has emerged (Ansari et al., 2012). In the last decade, the nascent field has built a bridge between classroom and laboratory to achieve critical insights on the dynamic neural processes involved in student learning (Davidesco et al., 2023; Dikker et al., 2017), drawing on theories and methodologies from cognitive neuroscience, education, and psychology in order to characterize the complex, dynamic process in naturalistic multimodal environments.

With physiological methods such as EEG recordings now feasible via portable and inexpensive devices, researchers can collect measures of cognitive engagement in real-world classrooms (Li, 2021). However, mobile EEG methods are prone to conceptual errors (i.e., over or misinterpreting results) and, in a technology-rich but experience-poor research environment, studies of attention and engagement require more holistic and time-tested methodologies to fill the gaps between current knowledge and open questions. Thankfully, decades of development and work have gone into classroom observation and student engagement improvement through video analysis of classroom dynamics (e.g., Pianta et al., 2012). As such, the rich knowledge gained through human (Goldberg et al., 2021) and machine (Aslan et al., 2019; Hur & Bosch, 2022) measurement strategies offers an opportunity to scaffold the growth of cognitive neuroscience into the classroom (Sümer et al., 2021).

Against this backdrop, we developed two new, research-based frameworks to achieve a more sophisticated, multidimensional understanding of the components of classroom engagement and their relationship with learning; we have given the two approaches, collectively, the handle *Present*, in reference to both the alert state of engaged students and the simultaneity of our measurements. The *Present* frameworks are derived from the Attention Relevance Confidence and Satisfaction model of motivational design (ARCS - Keller, 2012), which posits attention as the most proximal factor to new knowledge acquisition and, consequently, academic success. We implemented methods from the neuroscience laboratory in real-world classroom settings to chart a path towards a finer-grained understanding of the relationships between learning outcomes and affective, behavioral, and cognitive engagement. We leveraged the computational power of non-invasive technologies to collect simultaneous objective, subjective, and either neural or video-recorded measurements of engagement and learning outcomes, shifting from post-hoc evaluative methods to real-time, frequent, and concurrent assessments of classroom engagement.

The first framework (henceforth denoted **Study 1**) features a multimodal, multivariate approach that utilizes real-time EEG recordings of students to measure cognitive engagement, post-class subjective learning surveys to measure affective engagement, and in-class objective performance measures as a proxy for successful learning. The second framework (henceforth denoted **Study 2**) acts as a feasibility study, addressing some technical problems that arose in Study 1 and augmenting our approach by combining objective performance measures embedded in the lecture as a proxy for learning (per the ARCS model) with revised subjective measures to gauge classroom affective engagement and video recordings to capture behavioral engagement. Results indicate that both *Present* frameworks provide effective and comprehensive approaches to measuring multiple components of real-time classroom engagement, reduce technical and theoretical challenges of existing approaches, and hold promise for progress in educational research and practice.

## Materials & Procedure

### Protocol of Study 1

#### Participants

Eleven undergraduate students (mean age: 21.55 years ± 1.31 months; 7F, 4M; 3 sophomores, 5 juniors, 3 seniors; 5 Neuroscience and Behavior Majors; 4 Psychology Majors; 1 Cognitive Science Major, 1 Undeclared) from the Columbia University community enrolled in a six-week neuroscience course in the Summer of 2022 as part of their curricular program. Students agreed to participate in the study by signing a consent form before the beginning of data acquisition. Participation in the research activities was not mandatory, and students were able to opt out at any time. Over three-weeks of the course during active learning class periods, concurrent subjective measures of perceived learning and course engagement, objective content retention measures, electrophysiological (EEG) recordings, and video camera recordings were collected. Study protocol was reviewed and approved through the Columbia University Institutional Review Board.

#### Subjective Measure of Learning

Subjective reports of students’ feelings of their learning are increasingly being collected by researchers in order to examine the alignment between subjective experience and observable expression (Monkaresi et al., 2016). After each class period, we administered a Subjective Assessment of Learning Gained (SALG) in order to examine the affective component of engagement. The SALG takes between 3 to 5 minutes to complete and prompts students to provide a score (on a 7- point scale, wherein 1 is strongly disagree, scaling up to 7 as strongly agree) rating their agreement with statements that address their subjective feelings about their learning and engagement during class. The survey can be downloaded in the project repository, and a breakdown of each question is also available in the Supplementary Material section. The tool is derived from the Student Evaluation of Educational Quality (SEEQ) (Marsh, 1982), a widely used 33 question student feedback questionnaire with a robust factor structure, excellent reliability, and reasonable validity, and was expanded by our research team to include a total of 44 questions. We evaluated subjective student experience in class along the following sub-components: confidence (Qs 18 and 21), understanding (Qs 13-16, 24), self-valuation (Qs 10, 17, 26), course engagement (Qs 9 and 12), current study habits (Qs 11, 19, 23, 25), clarity of instruction (Qs 18, 22). The remaining questions queried students’ subjective feelings on particular instruction styles (e.g., active learning and lectures), and their academic demographics (e.g., planning to pursue a PhD, MA, etc.). Students’ responses were recorded after each class through Columbia University’s learning management system, Canvas. Each item was scored and subsequently fitted into one of six sub-components of emotional engagement. The means and standard deviations were then calculated to derive a subsequent ‘score’ for each sub-component, combining multiple items into one. For example, each student received a ‘subjective understanding’ score, based on their mean ratings of their responses to questions 13-16 and 24, for each class session.

#### Active Learning Measure

Student responses to content questions on Poll Everywhere, a web-based audience response system available on Android, iOS, and via browser apps, measured objective performance during course lectures. Poll Everywhere was freely available to participants and already used by one of the authors in his courses. Thus, students completed multiple choice quizzes (with either four or five answer choices) assessing content understanding and retention while it occurred, and they were able to self-evaluate their correctness through feedback provided by the instructor. All recorded measures of active learning were considered research activities, and didn’t hold any weight on students’ final grades. This measurement of content retention indexed each student’s retention of information within and across classes, granting students and instructors greater insight into learning as it unfolded while also affording more detailed characterization of engagement during a particular subject or instructional approach.

#### Mobile EEG

Several recent studies have used mobile electroencephalography (mEEG) to examine neural correlates of cognitive engagement (e.g., Bevilacqua et al., 2019; Halderman et al., 2021). In our study, Muse 2 portable EEG devices recorded students’ brain signals as a correlate of attention and engagement. Muse 2 (InteraXon, 2017) is an EEG device built into a headset worn across the user’s forehead and wrapped behind their ears. It has five sensors: two placed on a person’s forehead, two behind the ears, and a reference sensor positioned in the middle of the forehead (approximately FPz). Sensors on the forehead correspond to AF7 and AF8 electrodes in the 10-10 system, while the sensors behind the ears correspond to TP9 and TP10. Students were each assigned a Muse headset and corresponding laptop for signal streaming via Bluetooth lab streaming layers (muse-lsl) and a BLED112-V1 USB dongle. Source code was derived from Barachant, 2017 (https://github.com/urish/muse-lsl-python). Our code pipeline enables connection, streaming, live visualization, and marking EEG events, and can be found at the following link: https://github.com/clshatto/SOLER-Muse-EEG-Pipeline. Students were trained to fit the headset for optimal signal, and quality control was conducted by the research team before each recording session. Each student was also briefly taught how to visualize their own signal via the website EEGedu.com, which allowed the research team to review signal quality and assist in necessary adjustments to headset fit. However, students were not able to see their signal once quality checks were complete to ensure minimal disruption to the academic activities. Inspired by previous work utilizing Muse devices for brain signal detection (e.g., Krigolson et al., 2017, 2021), our protocol builds upon these findings to justify the adoption of Muse devices as the primary tool for detecting brain signals in a naturalistic classroom environment. By leveraging the expertise gained from both of Krigolson et al.’s studies, we aimed to employ the Muse technology to effectively assess cognitive performance and mental states, contributing to the understanding of real-world neurophysiological dynamics in educational contexts.

A dual-pass Butterworth filter with a passband of 0.1 to 50 Hz and a 60 Hz notch filter was applied to continuous EEG data. In order to eliminate sharp changes in signal most likely due to motion or eyeblink artifacts, we ran an independent components analysis to separate the remaining channels into independent components representing unique variance with respective time-series. The one component time-series that most resembled punctate motion or eyeblink artifacts, typically the first component, was then removed from the data. To increase our signal-to-noise ratio, we next created a pooled frontal and a pooled posterior virtual electrode by averaging across the frontal (AF7 and AF8) and the temporoparietal (TP9 and TP10) electrodes, respectively, as previously reported (Krigolson et al., 2021). Next, to eliminate discrete timepoints with still high fluctuations in signal, we divided the entire continuous data into 500 ms segments with 250 ms overlap, and segments that had an absolute difference of more than 60 microV were eliminated (on average: 33.3% [7.1%, 60.0%]). We used a Fast Fourier Transform to extract the power at various frequencies. For all statistical analyses, the absolute band power within the Theta range (4-8 Hz) was divided by the absolute band power within the Beta range (12-30 Hz) to find the average theta-beta ratio for each student across the duration of each class.

##### Statistical procedures

Average theta-beta ratios recorded during an entire class period, average active learning quiz performance, and average subjective learning responses for each student for each class period were related to each other with multilevel models using the lmerTest package (version = 3.1-3) in R (version = 4.0.3). Confidence, Understanding, Self-valuation, Course Engagement, Current Study Habits, and Clarity of Instruction scores were also used to examine the interrelationships between subjective scores and other components of engagement. To isolate the within-subject variance in our independent variables (theta-beta ratios and quiz performance), each student’s raw score from each class was subtracted from that student’s overall average score across every class. Each model included a random intercept for each subject crossed with a random intercept for each class, in order to separately account for variance across subjects and variance across classes, respectively.

To compare model predictive performance, models were submitted to the anova() function in the R stats package (version 4.0.3). This function calculates the likelihood-ratio between the models in order to compare the goodness of fit of each model to the data, assigning the deviance between them a value within the Chi-square distribution.

#### Video Camera Recordings

Two video cameras (Canon Vixia H R800) were placed in the classroom (**represented in Figure 1a)**. Due to space constraints, one camera was placed at a podium near the projection screen in front of a 12 x 4 feet table with students facing inward (**Figure 1b**). The other camera was placed on the table halfway down (**Figure 1c**).

**Figure 1.**
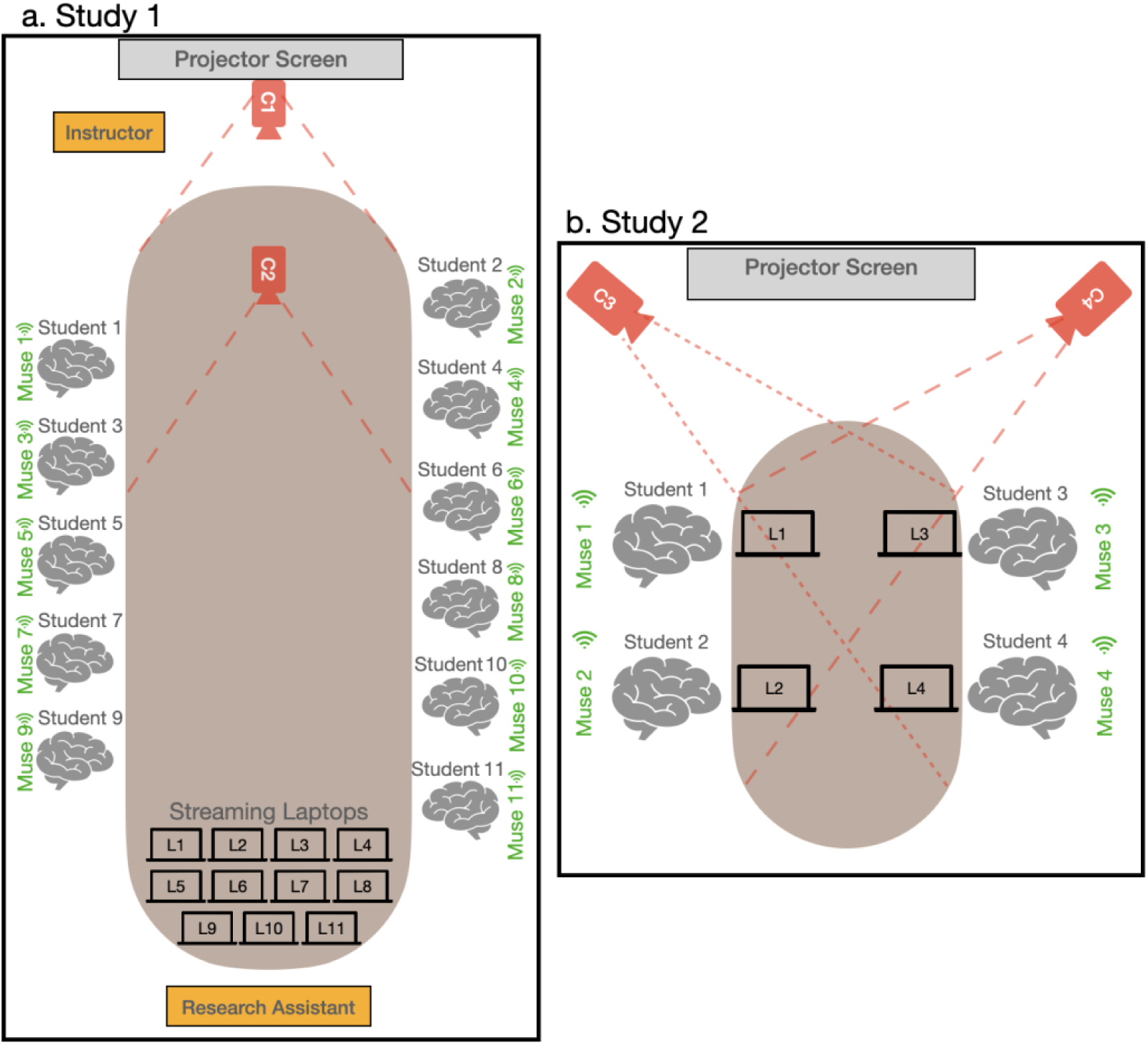
**a)** Schematic representation of classroom seating assignments in Study 1, showing camera positionings (c1 and c2). **b)** Schematic representation of classroom seating assignments in Study 2, showing the camera positionings for cameras c3 and c4.

##### Statistical Procedures

Data streams were initially assessed via open source facial quantification software (Baltrusaitis et al., 2018) and a custom manual coding system. We determined that the camera angles made the resulting data from both processes very unreliable.

### Results of Study 1

#### Active learning quiz performance is predictive of subjective understanding of course material

First, we were interested in the relationship between our measure of content retention and the SALG scores from the six sub-components. Multilevel models were separately run for each factor of the SALG predicting the average response for each student for each class session, with a fixed effect of within-student change in average quiz performance and random intercepts for each student and each class session. We found significant evidence that quiz performance was predictive of subsequent students’ subjective feelings of their understanding of course material (*B* = 0.73, t(12.9) = 2.27, *p* = .041) (**Figure 3a**). All other models reported little to no evidence of a relationship (all *p*s > .27). This indicates that, intuitively, students’ feeling of learning and objective learning scores were aligned, such that when a student performed better on the in-class quizzes than they usually did, they also reported higher subjective understanding of the material than they usually did.

**Figure 3.**
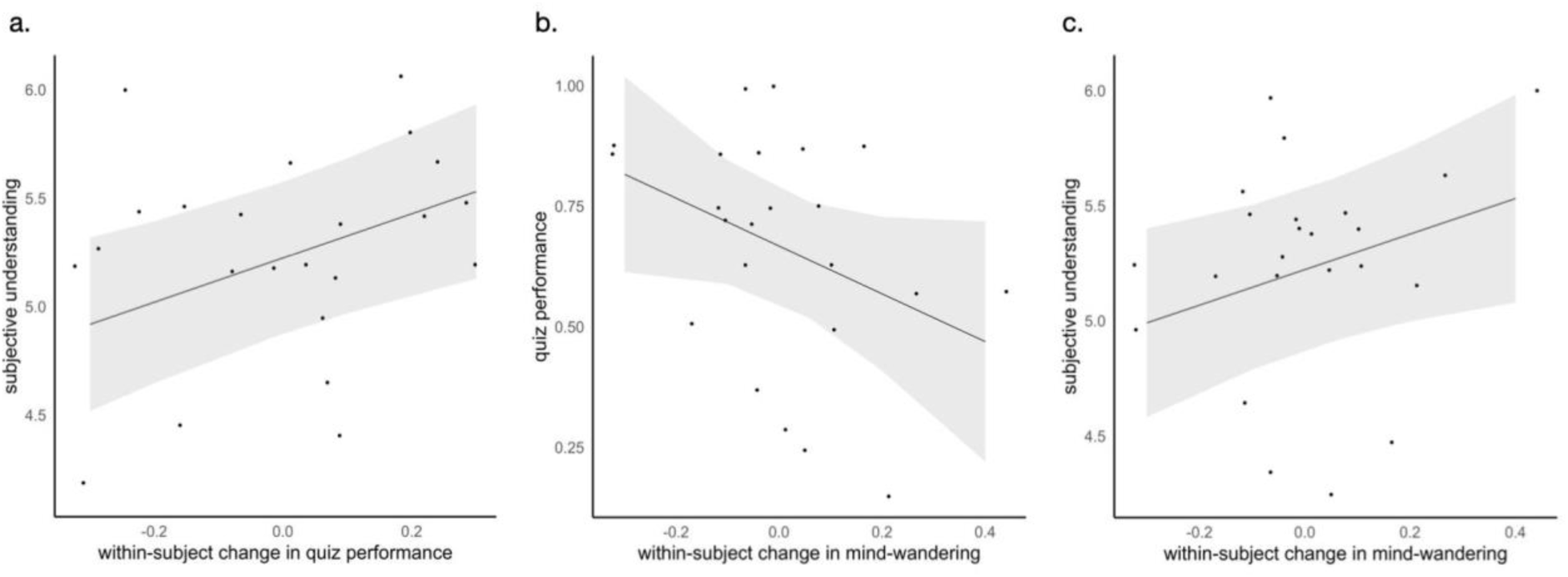
**a)** Objective quiz performance is positively predictive of subjective understanding. A multilevel model shows that within-subject change across classes in objective quiz performance is positively related to subsequent subjective feeling of understanding. **b)** Mind-wandering is marginally negatively predictive of quiz performance. A multilevel model shows that within-subject change across classes in theta/beta ratio, our neural metric of mind-wandering, is somewhat negatively related to in-class quiz performance. **c)** Mind-wandering and quiz performance together predict the subjective feeling of understanding material. A multilevel model shows that within-subject change across classes separately in theta/beta ratio and quiz performance are significantly related to subsequent ratings of subjective understanding of material.

#### Theta/Beta ratio is marginally negatively predictive of active learning performance

We next sought to characterize the relationship between our measures of mind-wandering during class, as indexed by the theta/beta ratio, and our measure of objective content retention. Theta/beta ratio has been previously confirmed to vary concurrently with mind-wandering states (Van Son et al., 2019), and attentional control (Department of Education and Child Studies, Leiden University, The Netherlands et al., 2020). Higher theta/beta ratios have been consistently observed during mind-wandering episodes, suggesting that this neurophysiological measure can serve as a reliable indicator of the occurrence and intensity of mind-wandering states. A multilevel model of the average quiz performance of each student for each class session was run, with a fixed effect of within-student change in theta/beta ratio and random intercepts for each student and each class session. We found marginal evidence that theta/beta ratios were negatively predictive of quiz performance (*B* = -0.50, t(14.1) = -1.86, *p* = .083) (**Figure 3b**). This suggests, intuitively, that when a student mind-wandered less during a class period, they performed better on the in-class quizzes.

#### Theta/Beta ratio alone is unpredictive of subjective feelings of learning

Next, we examined the relationship between theta/beta ratios and our measure of students’ subjective feelings of their learning. A multilevel model of the average subjective understanding rating for each student for each class session was run with a fixed effect of within-student change in theta/beta ratio and random intercepts for each student and each class session. We found no evidence that theta/beta ratios were predictive of subjective scores within the understanding sub-component (*B* = 0.34, t(12.8) = 0.83, *p* = .42) (**Figure 3c**). This indicates that, alone, a student’s mind-wandering during class will have little to no effect on how they rate their subjective feelings of their learning.

#### Multiple regressions of subjective understanding outperform individual regressions

So far, we have presented (1) evidence that better quiz performance leads to increases in subjective reports of content understanding and (2) marginal evidence that mind-wandering leads to poorer quiz performance. Taken alone, mind-wandering seems to have no effect on subsequent subjective understanding. In order to monitor all our variables simultaneously and establish a framework to involve as much of the classroom experience as possible, we entered within-subject change of both theta/beta ratio and quiz performance as multiple regressors of subjective understanding. Using pairwise likelihood-ratio tests, we found this multiple regression provided a better fit to the data (i.e., better predicted subjective understanding) than both theta/beta ratio (X^2^(1) = 8.92, p = .0028) and quiz performance (X^2^(1) = 5.19, p = .023) alone. Notably, effect estimates for both within-subject change in theta/beta ratio (*B* = 0.77, t(12.4) = 2.27, *p* = .043) (Figure 1) and within-subject change in quiz performance (*B* = 1.02, t(13.0) = 3.27, *p* = .007) (Figure 2) nearly doubled in size, indicating that they each accounted for separate sources of variance in the data.

**Figure 2.**
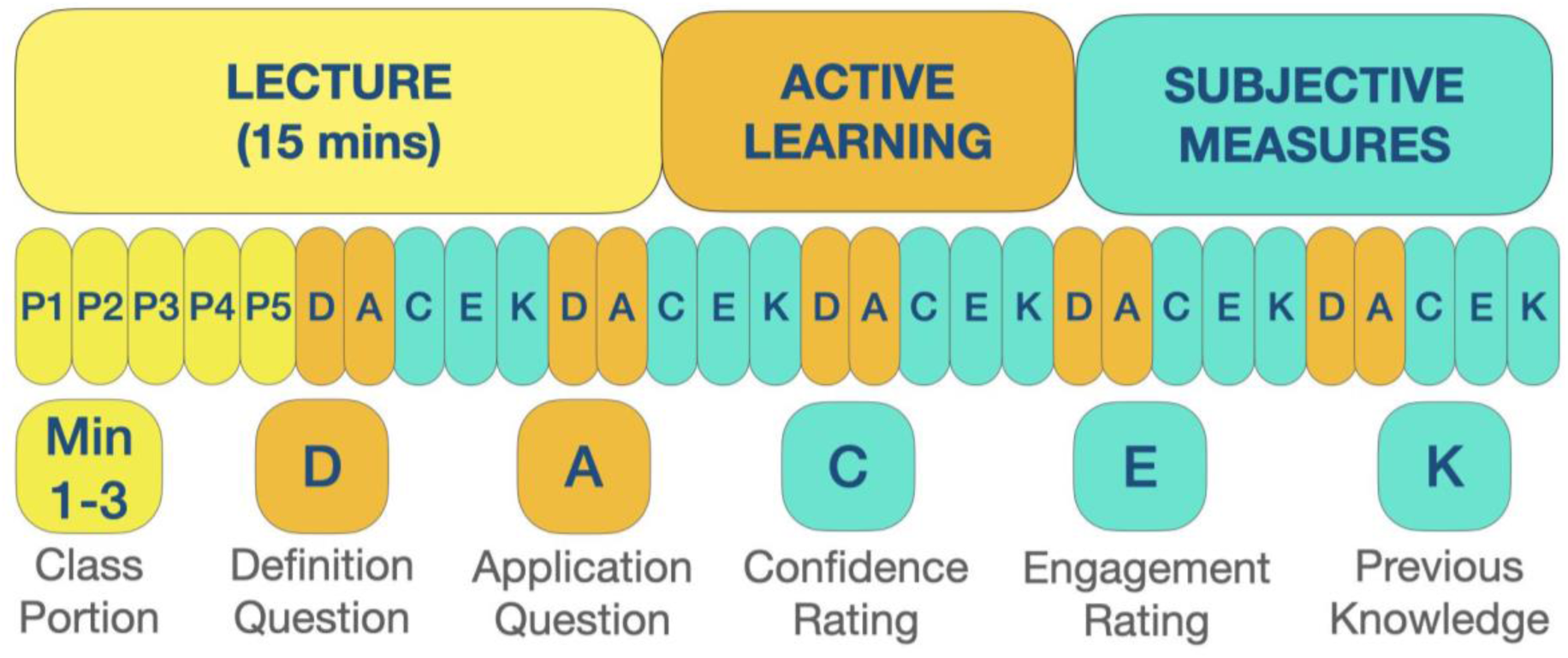
Schematic representation of classroom activities during Study 2. Participants watched two 15-minutes video lectures. Each lecture was followed by Definition (D) and Application (A) questions tapping on content retention, and Confidence (C), Engagement (E), and Pre-existing Knowledge (K) to measure subjective feelings of learning.

This pattern of results is in line with recent evidence that challenging, active participation enhanced subsequent quiz performance but weakened subjective feeling of learning, denoted across literature as the Dunning-Kruger Effect (Schlösser et al., 2013). In this way, our evidence indicates that our two measures – mind-wandering and quiz performance – affected subjective understanding most strongly when the other measure did not change. That is, mind-wandering most dramatically enhanced subjective understanding when quiz performance was consistent with other classes and, equivalently, quiz performance most dramatically enhanced subjective reports of understanding when mind-wandering was consistent with other classes.

#### Video Camera Recordings

Though intended to capture faces and body pose, our camera positionings resulted in students reciprocally obstructing the camera view at unpredictable intervals (e.g., students 4 and 10 in **Figure 1b**), which precluded reliable facial and pose tracking and standardization of estimation due to the variable depth and angles at which students were observed. Video camera measurements obtained under these camera positions were therefore excluded from analysis, and these obstacles informed our improved approach in Study 2.

### Interim Conclusion of Study 1

Study 1 indicated that objective quiz performance was predictive of students’ subjective understanding of course material. Further, examinations of classroom data revealed preliminary evidence that an EEG index of mind-wandering states —theta/beta ratio—is predictive of objective learning. These findings align with prior studies indicating that cognitive engagement is predictive of successful classroom learning (Greene et al., 2004; Rotgans & Schmidt, 2011). Mind-wandering states indexed by EEG theta/beta ratio were negatively correlated with quiz performance. This relationship was observed despite the numerous challenges associated with using consumer-grade mobile EEG devices. First, continuous EEG recordings are significantly limited by several factors: (1) acquisitions lasting more than 10-15 minutes often result in signal (packet) drops and loss of Bluetooth connection, likely due to technological limitations not identified by the manufacturer; (2) the presence of multiple Bluetooth devices streaming in one confined space (Study 1 attempted to collect data from 11 students in the same classroom) increases the time needed for stable Bluetooth connections to be established, likely due to radio frequency interferences; (3) device fit and wearability are inadequate over long durations, likely due to the device’s hard plastic construction and failure to achieve a tight fit over the forehead for the anterior-frontal sensors to make good contact; (4) the required one-to-one link between each device and a laptop due to the lab-streaming layer pipeline we used makes the technological setup more expensive in cost and needs – all must be individually connected and charged, a cumbersome arrangement (**Figure 1b, c**).

We have recently tested a new approach outside of a classroom setting, demonstrating the possibility to significantly reduce data loss from 30 seconds (in the case of manual Bluetooth reactivation performed by the RA as needed) to 5-6 seconds. Our code for the automatic Bluetooth reconnection can be found here: https://github.com/jnthorp/muse-lsl-cu. Alternative pipelines that allow linking multiple devices to one streaming layer already exist (e.g., BlueMuse, https://github.com/kowalej/BlueMuse). However, our attempts to record the signal using BlueMuse during our Study 2 protocol and synchronize it with markers related to the timing of in class quizzes were unsuccessful (discussed in further detail below). We believe that this issue will be overcome by future developments of streaming layer pipelines that will enable multiple device acquisition with seamlessly aligned markers.

The above-mentioned limitations motivated us to conduct a second study that addressed these obstacles in the following ways: (1) we limited our continuous data acquisition sessions to a maximum of 15 minutes; (2) we spaced out students within the same room in an attempt to reduce cross-device Bluetooth interference; (3) we used a newer version of the Muse device (Muse S), made of a softer fabric, to improve comfort over the course of an extended classroom session; (4) we modified camera positioning by adding tripods and placing the cameras approximately six feet away from the students and right next to the projector screen (as shown in **Figure 1d**) and zooming in on their faces. Additionally, we tested whether facial data could be recorded using student’s laptop cameras and examined the reliability and validity of our face detection algorithm on that data stream.

### Protocol of Study 2

#### Participants

Four individuals affiliated with Columbia University (mean age: 25.92 years ± 8.67 months; 2F, 2M; 2 undergraduates majoring in Neuroscience, 1 master’s student in clinical psychology, 1 PhD) participated in a single, 45-minute experimental learning session where they watched recorded lectures while being video recorded, and then answered post-test assessments of learning and engagement (**Figure 1 panel d**). All were asked to sign a consent form before the beginning of data acquisition. Participation in the research activities was voluntary, and all were able to opt out at any time. Study protocol was reviewed and approved through the Columbia University Institutional Review Board. One of the participants was an author of the manuscript, planted in order to test the impact of fluctuations in head and pose movements on video data acquisition.

#### Active Learning and Subjective Measures

We selected two 15-minute lectures from the YouTube TED Talk channel. At the end of each lecture, a post-test was conducted using Google Forms to obtain objective measures of learning via multiple-choice questions. Participants were first asked definition questions designed to identify a keyword (e.g., “[The Speaker] said he was a self-described_______”; **D in Figure 2**), and application questions designed to synthesize a theme in the lecture (e.g., “Big ideas, according to [The Speaker] require_______”; **A in Figure 2**). Following completion of the objective quiz questions, subjective measures of learning were collected via self-reported response confidence (**C in Figure 2**; rated on a scale of 1-5, with 1 being “not confident at all” and 5 being “absolutely confident”), level of engagement (**E in Figure 2**; rated on a scale of 1-5, 1 being “not engaged at all” and 5 being “absolutely engaged”), and whether previous knowledge was used to answer the D and A questions (**K in Figure 2**). Both types of questions were time-locked to particular portions of the video (i.e., questions D1, A1 required information provided in minutes 1-3 of the lecture, D2 and A2 from minutes 4-6, etc.). The five minutes after the post-test was spent debriefing participants, giving performance feedback, and providing a short break. This procedure allowed us to explore a novel measure of ground truth engagement (i.e., “Attend” below) defined by the combination of students’ responses to the objective retention questions and the presence of previous knowledge (**Table 1**), while maintaining the ability to analyze the relationship between subjective feelings and observed physical markers of attention and engagement during learning.

**Table 1.** Description of the “Attend” score estimated based on a combination of content retention questions and previous knowledge checks.

| “Attend” score | Previous Knowledge | Definition Question | Application Question |
| --- | --- | --- | --- |
| Drop | Yes |  |  |
| Guess | I don’t know | Correct \| Incorrect | Correct \| Incorrect |
| Guess | I don’t know | Correct \| Incorrect | Correct \| Incorrect |
| 0 | No | Incorrect |  |
| 0 | No | Correct | Incorrect |
| 1 | No | Correct | Correct |

#### EEG Recordings

Here, Muse S portable EEG devices recorded students’ brain signals correlated with students’ attentional focus and engagement. Muse S (InteraXon, 2017) is an EEG device built into an elastic headband worn across the student’s forehead. Similarly to Muse 2 (Study 1), it has five sensors: two placed on a person’s forehead, two behind the ears, and a reference sensor positioned in the middle of the forehead (approximately FPz). Sensors on the forehead correspond to AF7 and AF8 electrodes in the 10-10 system, whereas the sensors behind the ears correspond to TP9 and TP10. Streaming and recording were conducted using the same procedures as in Study 1.

#### Video Camera Recordings

Two cameras at the front of the classroom and built-in laptop cameras were utilized to record overt behaviors related to attentional focus, such as student gaze tracking, head pose, and facial muscle movements. Each camera at the front was positioned to capture two participants on either side of the room from above, allowing a clear view of the face and upper body of each participant. This setup was chosen to correct problems with video camera data acquisition encountered in Study 1. **Figure 1d** shows a schematic of the classroom setup. Videos were trimmed to include only the period when the TED Talk was playing and cropped to include only one participant in the field of view via a custom ffmpeg script.

Each video was then processed as follows: (1) head pose, face segmentation, facial landmarking, gaze direction, and facial action unit (FAU) presence and activation amplitudes were estimated via OpenFace 2.0 (Baltrusaitis et al., 2018); (2) processed videos were analyzed for reliability of facial landmark tracking through visual analysis of outputs, and confidence and success ratings from OpenFace; (3) all videos that were deemed unreliable were excluded from further analysis; (4) all feature vectors of reliable videos were smoothed with a Gaussian filter (Nadaraya, 1964) with an optimally chosen bandwidth according to Wand and Jones (1994) to optimize signal to noise ratios; (5) objective and subjective measures were merged with each participants’ face data by assigning each frame a ‘1’ for definition (D), application (A), and previous knowledge (K) questions if correct (or answered based on previous knowledge; ‘0’ if incorrect or not based on previous knowledge) and the self-reported rating (1-5) for both confidence (C) and engagement (E) measures.

Overall, this process identified four videos from two participants (one for each TED Talk) reliable enough for further analysis; the other two participants’ videos were excluded. One participant was excluded due to wearing a facemask for both recordings. The other participant (see student 3 in **Figure 1e, 1f**) was excluded because they spent about half of the video moving, looking away from the camera, putting their hands in front of their face, and drinking from a water bottle. Note that this participant was asked to move frequently on purpose to estimate the algorithms’ performance under suboptimal conditions (i.e., by simulating a ‘restless’ student rather than one staying still). Laptop videos were also excluded from data analysis because they captured only the profile view of each participant. Efforts to address these known issues constitute our internal feasibility case study (described in more detail below).

#### Statistical procedures

Smoothed and labeled facial data were used for three separate but related exploratory analyses: (1) an analysis of correlations between all variables of interest, (2) linear regression analyses with the lm() and glm() functions in R predicting subjective (continuous) and objective (binomial) performance respectively with facial data across individuals and videos, and (3) a machine learning analysis predicting ‘Attend’ defined as ‘1’ if both D and A were responded to correctly, and if the participant’s answer was not based on previous knowledge, and ‘0’ if any of these conditions were not met (**Table 1**). Overall, we predicted that head pose and gaze direction variables would be most associated with and predictive of attention, and AU04 (Brow Lowerer) and AU07 (Lid Tightener) would be related to attention, but spuriously so in the linear models. We also predicted that adding individualizing information into the dataset would improve AUC metrics in the machine learning models, as would a radial, as opposed to linear, kernel. For the correlational and linear regression analyses, we calculated the average amplitudes of every facial dimension individually within every question for each participant. For linear regressions, the anova() function in R was used as in Study 1 to compare and evaluate models. For the machine learning analysis, we randomly split the dataset into training (75%) and test (25%) components.

### Results of Study 2

#### Reliability of the video camera data depends heavily on camera angle and facial obstruction

Results of our internal feasibility study suggest that video camera data collection is hindered by restless movement in participants and is unreliable in the presence of face masks when applying OpenFace 2.0 to classroom video data. Videos of participants who were naturally watching the TED talks without face masks on showed very high levels of reliability (*μ* = .97, *σ* = .02), whereas our restless participant’s videos were lower (*μ* = .90, *σ* = .16), and our face mask participant’s videos even lower (*μ* = .81, *σ* = .34; see **Figure 4**). All participants’ laptop videos showed lower reliability (*μ* = .71, *σ* = .37) and often led to unsatisfactory results even when rated with high confidence by the algorithm (see **Figure 5**). These results indicate the necessity of caution when analyzing facial data through automated means and justify our feasibility case study to evaluate proper methodology for this type of data collection. Notably, during the second TED talk (in comparison to the first) participants were wearing Muse2 headbands to record EEG data, and the headband covering a large portion of their forehead did not appear to lower the quality of the facial data collection by OpenFace

**Figure 4.**
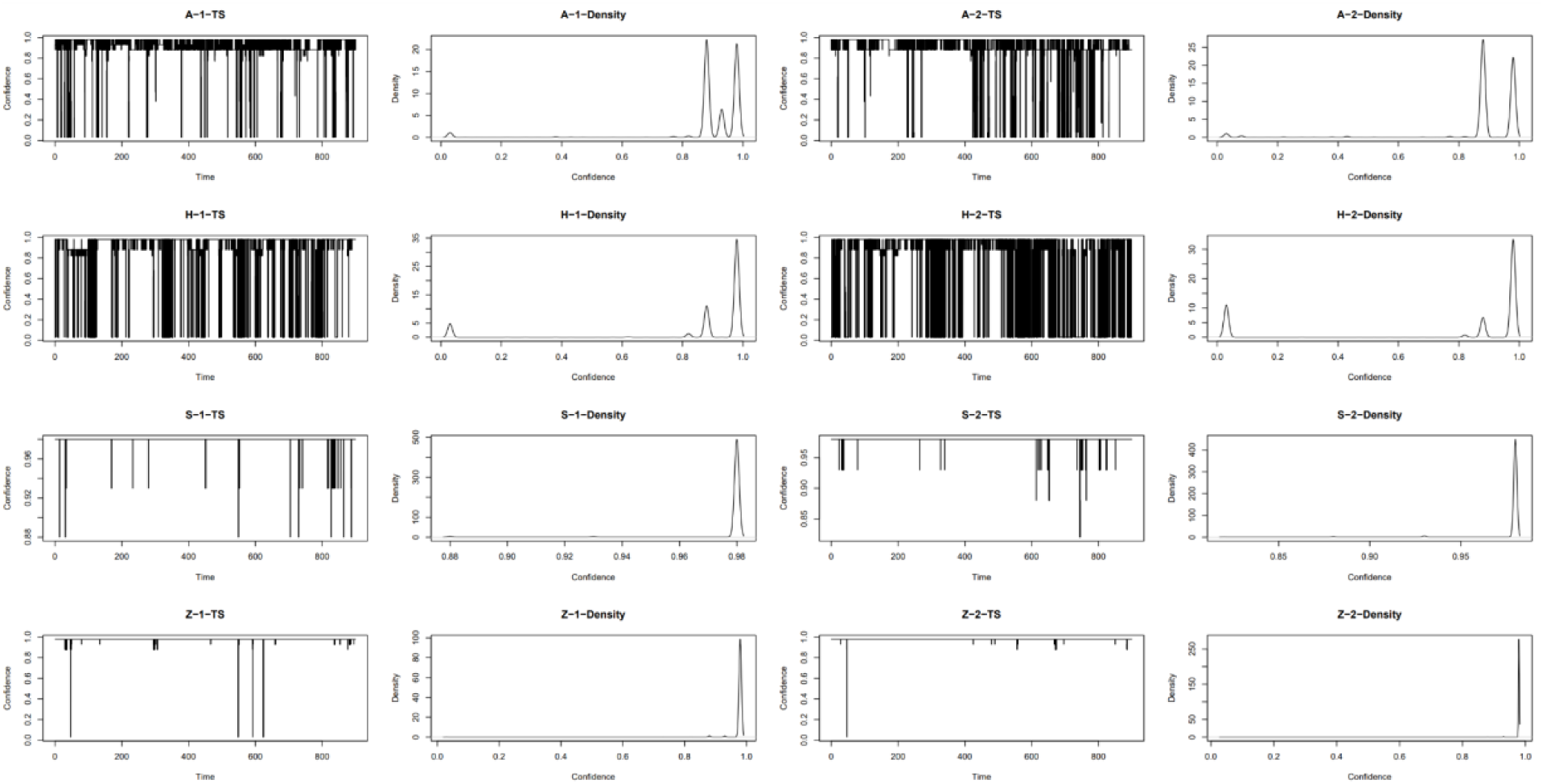
OpenFace 2.0 reliability measures (i.e., ‘confidence’) for all Camera videos in Study 2 in time series (TS) and Density plots organized by participant and video.

**Figure 5.**
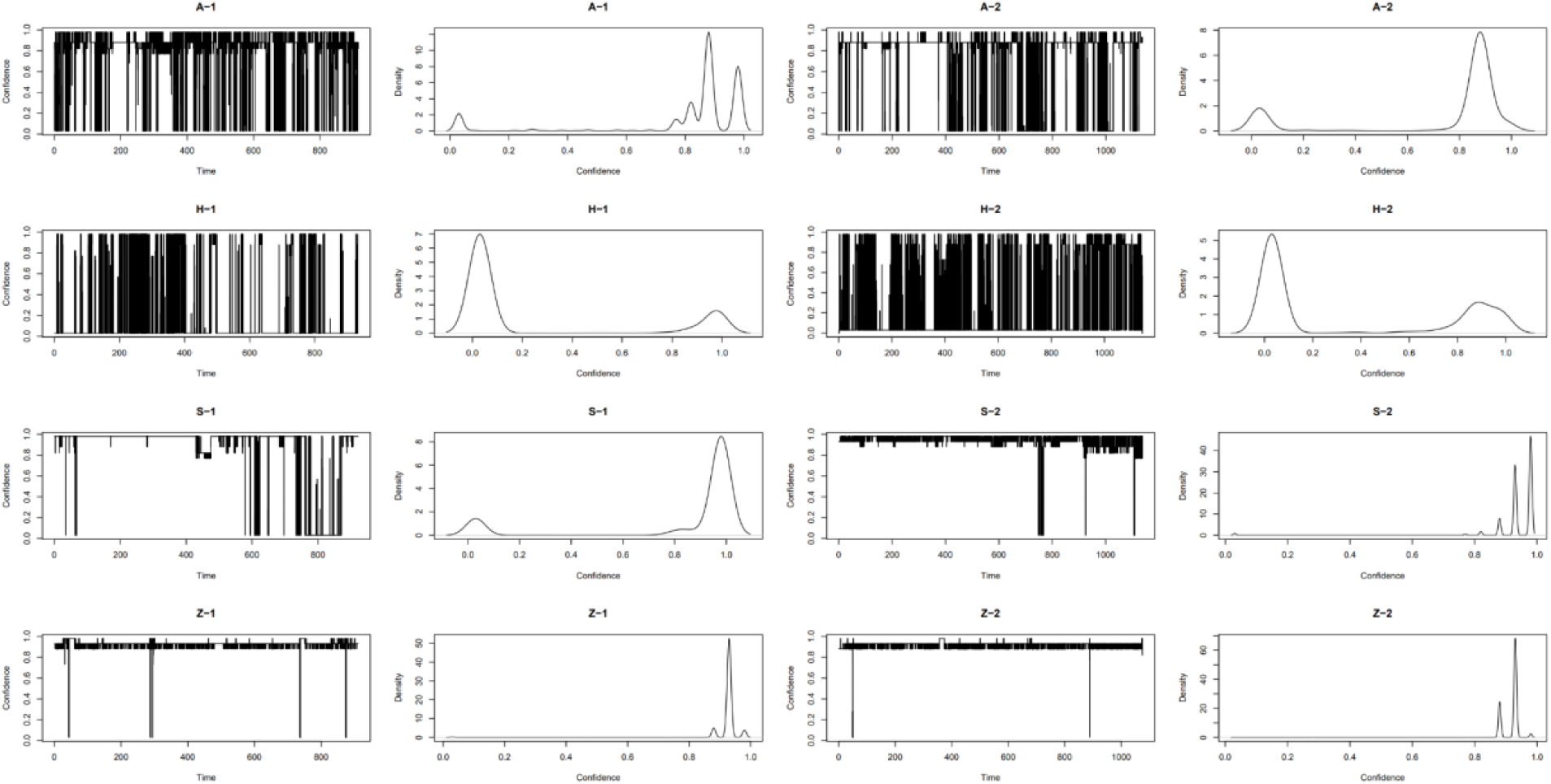
OpenFace 2.0 reliability measures (i.e., ‘confidence’) for all Laptop videos in Study 2 in time series and Density plots organized by participant and video.

2.0. After this reliability analysis, we included all 215,707 frames of video data from 2 participants across 2 videos who were ‘naturally viewing’ the TED talks for the analyses below. Based on the obstacles encountered during video camera analyses in Study 1, we adjusted our camera placement strategy for Study 2 (**Figure 1d**), and the high rates of tracking and estimation successes attained in Study 2 underscore the need to intentionally design camera angles and use caution when collecting, analyzing, and interpreting face and pose data from video recordings ‘in the wild’.

#### Single dimensions of facial data show reliable associations with subjective, but not objective measures

Among 22 facial data variables, none was significantly correlated with definition or application responses (all *p’*s > .12; see **Figure 6**). However, confidence was associated with gaze (*r_GazeX_* = -.51; *p_GazeX_* = .02; *r_GazeY_* = -.61; *p_GazeY_* = .004) and head pose (*r_Yaw_* = -.58; *p_Yaw_* = .006; *r_Pitch_* = .48; *p_Pitch_* = .03), as well as AU04 (*r* = -.6; *p* = .004) and AU07 (*r* = -.51; *p* = .02). Engagement was associated with gaze (*r_GazeY_* = -.52; *p_GazeY_* = .01), head pose (*r_Yaw_* = -.47; *p_Yaw_* = .04), AU04 (*r* = -.53; *p* = .02) and AU10 (*r* = .52; *p* = .02).

**Figure 6.**
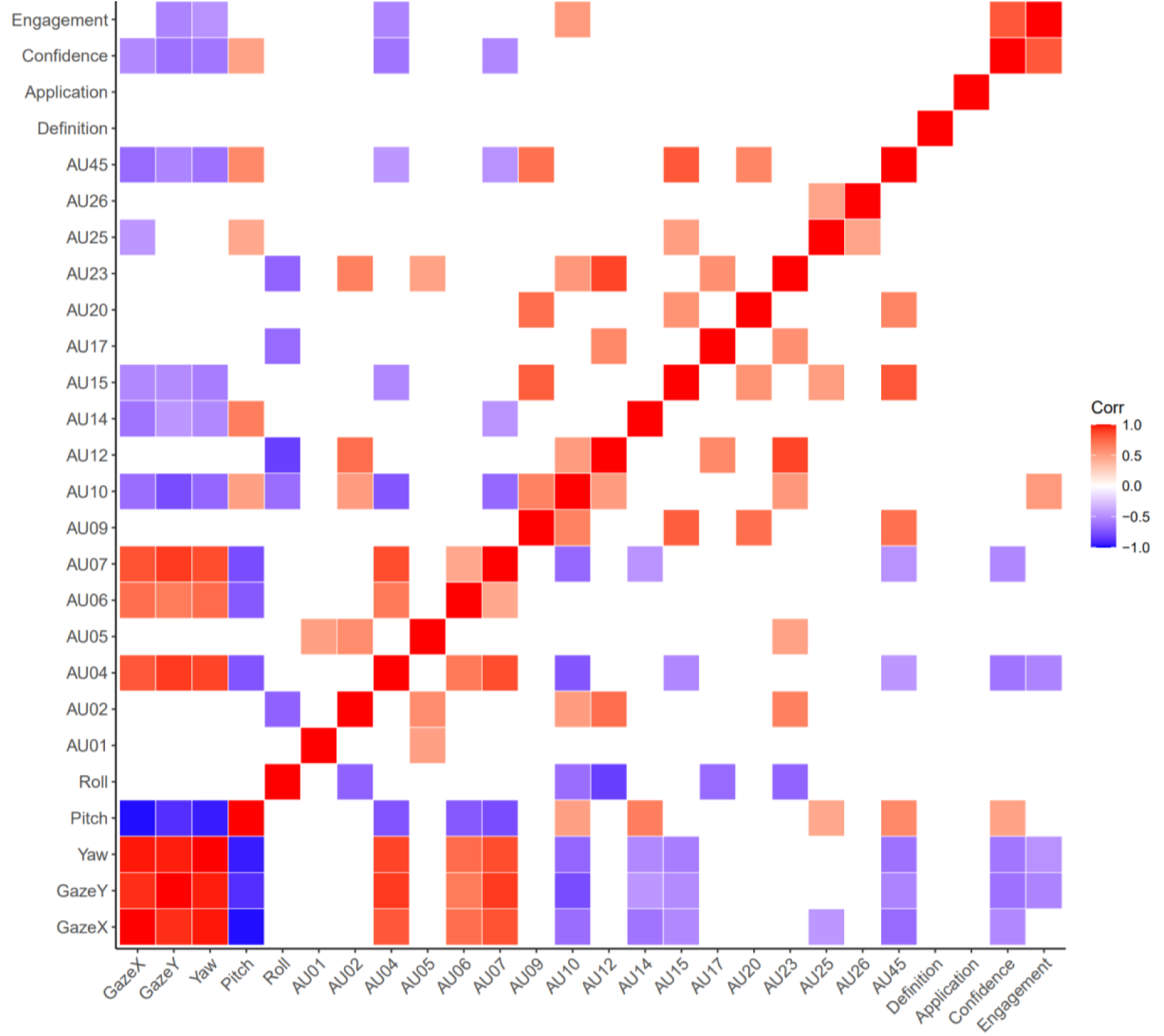
Bivariate correlations between facial data, subjective, and objective performance Measures. Only correlations with p-values < .05 are shown.

#### Both subjective and objective measures can be predicted by multiple dimensions of facial data

Yaw and Pitch of the head significantly predicted performance on application questions (LR *χ^2^* = 9.77; *p_Yaw_* = .001; LR *χ^2^_pitch_*= 9.2; *p_pitch_*= .002), but not on definition questions (all *p*’s > .2). Neither roll of the head, nor gaze direction predicted objective performance measures (all *p*’s > .2), and neither did AU04, AU07, AU10, or a model with any combination of them (all *p*’s > .14). Subjective measures, on the other hand, were predicted by head pose, gaze direction, and FAU amplitudes. Engagement was significantly predicted by: (1) head pose (*F*_2,17_ = 5.76; *p* = .01; *r*^2^ = .33), specifically by Yaw (*t* = -2.52; *p* = .02) and Roll (*t* = -2.27; *p* = .04); (2) gaze direction (*F*_2,17_ = 4.92; *p* = .02; *r*^2^ = .29), specifically by the Y dimension (*t* = -2.38; *p* = .03) but not X (*t* = -1.56; *p* = .13); and (3) FAU amplitudes in separate models (AU04 - Brow Lowerer: *F*_1,18_ = 7.13; *p* = .02; *r*^2^ = .28; AU10 - Upper Lip Raiser: *F*_1,18_ = 6.63; *p* = .02; *r*^2^ = .27). Confidence was significantly predicted by: (1) head pose in separate models (Yaw: *F*_1,18_ = 9.43; *p* < .01; *r*^2^ = .34; Pitch: *F*_1,18_ = 5.492; *p* = .03; *r*^2^ = .23); (2) gaze direction (*F*_2,17_ = 6.14; *p* < .01; *r*^2^*_adj_* = .35), specifically by the Y dimension (*t* = -2.18; *p* = .04) but not X (*t* = 1.14; *p* = .26); and (3) FAU amplitudes in separate models (AU04: *F*_1,18_ = 10.25; *p* < .01; *r*^2^ = .36; AU07 - Lid Tightener: *F*_1,18_ = 6.61; *p* = .02; *r*^2^ = .27).

Interestingly, in most cases, single predictors of hypothesized relationships were consistently predictive of subjective measures, while models with multiple predictors sometimes predicted subjective and objective outcomes at greater than chance levels, but less often. Considering the categories of head, gaze, and facial action data, multicollinearity only became problematic when predictors across categories were all included in the same model (e.g., Gaze X, Pitch, and AU07; all VIFs > 9), but not when multiple within-category predictors were tested together (e.g., Pitch, Yaw, and Roll of the head; all VIFs < 5). Nonetheless, it remains possible that multiple across category predictors may enhance the predictability of the models, but our case study data does not possess enough power to detect it. These results suggest either artifacts from the OpenFace algorithm that uses pose direction in its calculation of facial landmarks, and facial landmarks in its calculation of gaze direction, or an inherent association between head pose and eye gaze direction during attention tasks (such as watching a video lecture). Finally, it is encouraging to note that using our “Attend” measure (as described in Statistical Procedures for Study 2, and in **Table 1**), we achieved similar results to predicting D and A performance alone, with the exception that Gaze Y was also a marginally significant predictor of Attending (LR *χ^2^*= 3.32; *p* = .07), as was AU10 (LR *χ^2^* = 2.81; *p* = .09) in within-category binomial models.

#### Individualization and radial kernels improve AUC metrics for machine learning models of Attention

Finally, in an attempt to overcome the multicollinearity issues inherent in the facial data while still extracting useful information from our feasibility study, we trained multiple classifiers with support vector machines (SVMs) using the svm() function in the e1071 package in R to predict “Attend” (see **Table 1**), incorporating all facial data and either (a) individualization among subjects, (b) radial kernels, (c) both of these, or (d) neither of these (see results in Figure 7). Interestingly, the radial model without individualization (i.e., a tag on every frame with the participant’s ID) performed best (AUC = .999), but only slightly better than with individualization included (AUC = .998), though this difference was significant (*D_98612_* = 3.62; *p* = .0002; DeLong et al., 1988) and this may be due to having only two participants sitting at very similar angles to the camera. The linear classifiers both performed much worse than the radial classifiers (all *p*’s < .0001). The linear classifier without ID labels also performed slightly better (AUC = .858) than the classifier with ID labels (AUC = .857), but this difference was not significant (*D_107850_* = 0.33; *p* = .74). Overall, these results suggest a radial multivariate distribution of facial data’s ability to predict our Attend measure in our limited and likely overfitted model, wherein there is a ‘goldilocks’ zone of facial configuration (not too activated or stretched, and not too relaxed or neutral) that indicates whether or not someone is attending, in this case, to a TED Talk.

**Figure 7:**
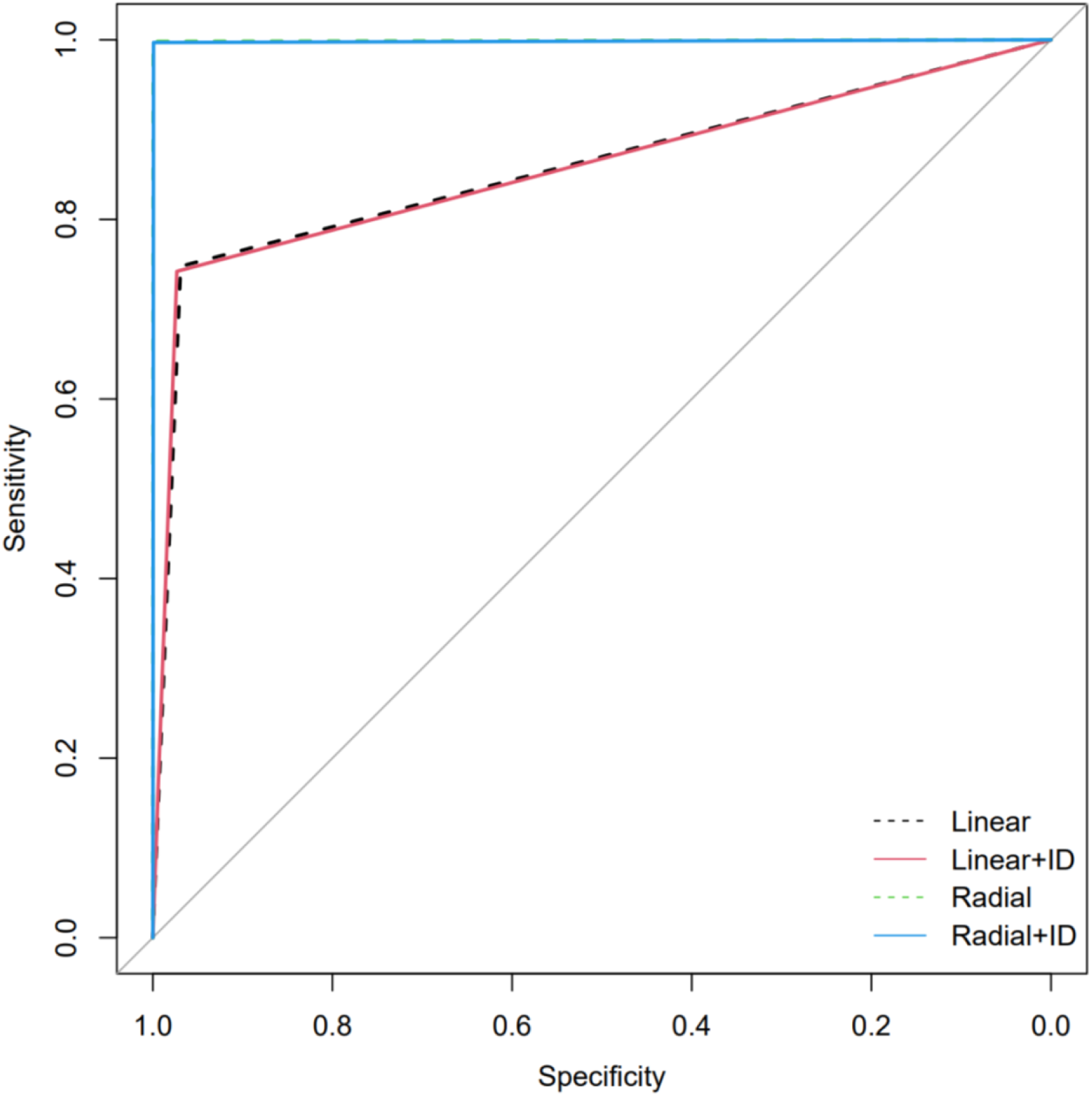
ROC Curves for Linear, Linear w/ Identification, Radial, and Radial w/ Identification SVM classifiers predicting “Attend” measure across 2 participants and 2 TED Talks.

#### EEG Recordings

Though intended to measure brain wave activity within a naturalistic classroom environment, these recordings were unsuccessful due to many drops in Bluetooth signal. Therefore, EEG measurements obtained were not further analyzed in our results. We discuss alternative solutions to these challenges below.

### Interim Conclusion of Study 2

Results from Study 2 indicated that subjective measures of engagement and confidence were positively correlated; contrary to our findings in Study 1, however, we did not observe any significant relationship between objective and subjective measures of learning. We also found that facial data can be utilized to predict students’ subjective feelings of learning and their quiz performance. Although this relationship was observed in a small sample size, our method shows promise for implementation in larger classroom environments.

In stark contrast with EEG data collected in Study 1, where challenges lay mainly in the scant progress in data acquisition strategies associated with new technology, obstacles affecting continuous video acquisition in Study 2 concerned achieving multiple camera angles for face and pose detection. Our framework prioritized ecological validity, collecting as much data as possible without compromising naturally occurring classroom behavior and dynamics. Hence, the first option to record continuous close-ups video of students’ faces was to use the built-in cameras on students’ class laptops. Below, we discuss challenges identified in our exploration of camera placement and potential solutions for these issues:

1. Laptops are usually placed on the classroom table, but students’ faces and gaze are most often directed towards the projector screen and the instructor; as such, camera angle becomes perpendicular to students faces and gaze (see **Figure 1f**), which precludes robust and reliable detection. Moving laptop cameras to fall in line between the students’ face and gaze and the projector screen could correct camera angle issues.
2. Utilizing overhead camera positioning improved face and pose detection data from each student, as discussed in the results section. However, our setup was a particularly favorable one, with two cameras used to detect only two students on each side of the classroom table; we expect this ratio to be the most reliable, which poses a challenge to studies interested in scaling up our approach to classrooms of 20 or more students. In Study 2, we used professional cameras on tripods to record students’ faces, but it is hard to imagine a class with a 2:1 ratio of students to cameras. Further, these professional cameras feature large objective lenses and are cumbersome, potentially compromising the ecological validity of the learning setting. Smaller, less conspicuous cameras could be more easily placed in the classroom at an angle that is favorable for data acquisition.
3. Masks, glasses with corrective lenses, and other facial features (e.g., beard) disrupted the face and pose detection algorithm regardless of camera placement. Little can be done about this, so researchers attempting to use our approaches must be aware of missing potential datapoints.

We were partly able to address another technical problem, EEG signal drops, by reducing interference between signal streams and increasing spacing between students. Given that these issues largely persisted, however, we believe a custom code that automatically re-establishes Bluetooth connection is necessary.

Overall, Study 2 succeeded in identifying better camera positioning for reliable collection and analysis of facial data and embedding objective and subjective measures of learning seamlessly within a 15-minute lecture. Below, we discuss how, taken together, the results from our two studies pave the way towards combining two technological and non-invasive, research-based frameworks to characterize classroom engagement.

## General Discussion

Understanding the dynamic nature of engagement in the classroom has important implications for learning and academic achievement. Broadly, we described two approaches that help set a foundation for concurrent evaluations of the components of classroom engagement, with minimal disruption to regular classroom activities and learning process. We further detailed the feasibility and utility of collecting EEG or video camera recording data within a naturalistic learning environment, such as an undergraduate level course, and related those data streams to subjective and objective measures of learning. Below we summarize four main takeaways from our studies and discuss how they can be used to guide future efforts in naturalistic studies of classroom attention.

First, we observed a positive relationship between successful learning and indices of affective engagement. This result supports what previous studies (Fredricks & McColskey, 2012) and many instructors already report: content retention is strongly related to students’ emotional reactions and feelings towards their learning in the classroom. Additionally, our results provide evidence that this objective-subjective association is strong enough to be measured asynchronously. In Study 1, affective engagement was measured by post-class subjective surveys whereas successful learning was measured in class. By including post-class subjective surveys of affective engagement to measure students’ interest, enjoyment, and enthusiasm, our methodology enables a more detailed understanding of successful learning. Interestingly, we were not able to replicate this finding in our Study 2 protocol, perhaps due to the reduction of items in our subjective evaluations of affective engagement. In Study 1, we gathered data from 33 items across four sessions with a 7-point scale to assess students’ subjective feelings about their learning. In contrast, Study 2 employed a condensed four-item query of students’ confidence and engagement administered only once, after a 15-minute learning period. Study 2 therefore allowed for a more time-efficient evaluation while still capturing key aspects of students’ affective experiences during the learning process, but may not have given participants the time and practice required to evaluate their engagement precisely. We are confident that collecting measures of affective engagement over multiple sessions throughout a course holds the potential to pinpoint the association between feelings of learning and actual learning more conclusively than we did in either study reported here.

Second, we observed a negative relationship between successful learning (as measured by in-class polls) and mind-wandering states in the classroom. In Study 1, cognitive engagement was indexed by EEG recordings through changes in theta/beta ratios, an established proxy for mind-wandering states. In line with previous studies, we showed that fluctuations in theta/beta ratios were associated with performance on in-class polls. We successfully implemented this methodology in the classroom, thereby uncovering covert processes of learning and cognitive strategies that are highly predictive of successful learning (Greene et al., 2004). Like our observed association between video recordings of in-class activities and successful learning, the association between cognitive engagement and content retention aligns with the ARCS framework. By including these brain-based measurements of cognitive engagement into evaluations of student learning, we can achieve a more comprehensive understanding of successful learning in the classroom, and better predict it to evaluate course and lesson design principles.

Third, we observed a positive relationship between successful learning measured by in-class polls and behavioral indices of classroom engagement collected via visual recordings. The relevance of this relationship is twofold. First, it provides scientific evidence supporting what experienced instructors already report: overt behaviors and visual indicators of students on-task or actively participating are meaningful barometers of student learning. Second, the observed relationship directly links behavioral indices of classroom engagement – namely, measurements of specific head movements – to accuracy on content application questions. Although certainly underpowered (as noted above), this result aligns with the ARCS framework. As such, automated, visual recording-based measurements of behavioral engagement constitute another powerful tool for characterizing and predicting classroom learning under appropriate circumstances.

Lastly, we observed a positive relationship between indices of affective and behavioral engagement. Like the preceding result, this finding empirically reinforces another principle of teaching that instructors intuitively understand: observable in-class behaviors such as paying attention and participating in activities are associated with students’ emotions surrounding learning. Furthermore, this result is novel because it links physical, behavioral indicators of students’ engagement to their *self-reported* (i.e., survey-based) affective engagement. Taken together, these results indicate a learning space for educators to better capture successful learning by connecting overtly observable classroom behaviors to students’ motivation, interest, enjoyment, and enthusiasm.

Overall, our two studies replicated prior findings of associations between each component of engagement and successful learning while also providing preliminary evidence for the association between behavioral and affective engagement. Together, our results demonstrate the feasibility and utility of bringing neuroscientific and computational methods into the classroom setting, with the potential to revolutionize current teaching and learning practices. Despite the challenges of bringing neuroscience into the “real world”, we believe that combining the approaches of our two studies can achieve a real-time characterization of the components of classroom engagement and their predictive relationships with successful learning. We presented a path to develop an advanced, automated assessment of student engagement, and we showed how we implemented this into a neuroscience seminar course at Columbia University. Bringing the neuroscience laboratory into the classroom is possible, but it can only be achieved by carefully aligning teaching and research activities. We showed that student learning can be assessed during 15-minute lecture portions via in-class quizzes (objective measures of learning), surveys about their response confidence (subjective judgments of learning), EEG recordings via lightweight, mobile headsets, and video recordings of face and pose expressions as well as large classroom interactions (via overhead cameras). We hypothesize that the relative value (i.e., explained variance) of each data stream can be used to predict engagement at the individual student level, and using an unsupervised classification model to reduce data dimensionality will help in this effort, likely in unexpected ways.

As in all settings, engagement during class fluctuates naturally due to physical, emotional, and social preoccupations. Distractions can be especially detrimental in STEM learning contexts, which are highly cumulative, information-dense and problem solving-focused: a momentary lapse may be compounded, as failure to attend to one concept may prevent understanding what follows even if focus is restored. Our approaches aim to advance our understanding of classroom engagement by quantifying the relative and combined contributions of the subcomponents of classroom engagement, so that they can be used as a tool by instructors to understand their students. Combining neuroscience principles and related technology with elements of the scholarship of teaching and learning, ‘Present’ constitutes a major step toward developing a rigorous, quantitative framework for investigating student experience in a variety of classroom settings.

## Supplementary Material

## SALG BREAKDOWN: SUBCOMPONENT SCORES

### Subcomponent 1) Confidence: Q 18 & Q 21

**Q18: Presently, I am confident that I understand the foundations of neuroscience.**

0: Not Applicable; 1: Strongly disagree; 2: Disagree; 3: More or less disagree; 4 Undecided; 5 More or less agree; 6: Agree; 7: Strongly Agree

**Q21: Presently, I am confident that I can be successful in this course.**

0: Not Applicable; 1: Strongly disagree; 2: Disagree; 3: More or less disagree; 4 Undecided; 5 More or less agree; 6: Agree; 7: Strongly Agree

### Subcomponent 2) Understanding of Material: Qs 13-16, 24

**Q 13. Presently, I understand neuroanatomy and physiology (how the anatomy of the brain relates to function).**

0: Not Applicable; 1: Strongly disagree; 2: Disagree; 3: More or less disagree; 4 Undecided; 5 More or less agree; 6: Agree; 7: Strongly Agree

**Q 14. Presently, I understand neuron morphology and physiology (how neurons ’fire’ action potentials).**

0: Not Applicable; 1: Strongly disagree; 2: Disagree; 3: More or less disagree; 4 Undecided; 5 More or less agree; 6: Agree; 7: Strongly Agree

**Q 15. Presently, I understand how the brain encodes our experiences of the world.**

0: Not Applicable; 1: Strongly disagree; 2: Disagree; 3: More or less disagree; 4 Undecided; 5 More or less agree; 6: Agree; 7: Strongly Agree

**Q 16. Presently, I understand how ideas we will explore in this class relate to ideas I have encountered in classes outside of this subject area.**

0: Not Applicable; 1: Strongly disagree; 2: Disagree; 3: More or less disagree; 4 Undecided; 5 More or less agree; 6: Agree; 7: Strongly Agree

**Q 24. Presently I understand the different theories of consciousness.**

0: Not Applicable; 1: Strongly disagree; 2: Disagree; 3: More or less disagree; 4 Undecided; 5 More or less agree; 6: Agree; 7: Strongly Agree

### Subcomponent 3) Self Evaluation of Skill: Qs 10, 17, 26

**Q10. Presently, I can critically read research articles about issues raised in class.**

0: Not Applicable; 1: Strongly disagree; 2: Disagree; 3: More or less disagree; 4 Undecided; 5 More or less agree; 6: Agree; 7: Strongly Agree

**Q17. Presently, I can share good ideas in discussion with the class.**

0: Not Applicable; 1: Strongly disagree; 2: Disagree; 3: More or less disagree; 4 Undecided; 5 More or less agree; 6: Agree; 7: Strongly Agree

**Q26. Presently I can share good ideas in lab meetings.**

0: Not Applicable; 1: Strongly disagree; 2: Disagree; 3: More or less disagree; 4 Undecided; 5 More or less agree; 6: Agree; 7: Strongly Agree

### Subcomponent 4) Course Engagement: Qs 9, 12

**Q 9. I study in groups.**

Not Applicable - 0 Strongly disagree - 1 Disagree - 2 More or less disagree - 3 Undecided - 4 More or less agree - 5 Agree - 6 Strongly agree - 7

**Q 12. I ask for clarification from my professor when I have questions.**

Not Applicable - 0 Strongly disagree - 1 Disagree - 2 More or less disagree - 3 Undecided - 4 More or less agree - 5 Agree - 6 Strongly agree - 7

### Subcomponent 5) Current Study Habits: 11, 19, 23, 25

**Q11. I am in the habit of sharing what I learn with others.**

Not Applicable - 0 Strongly disagree - 1 Disagree - 2 More or less disagree - 3 Undecided - 4 More or less agree - 5 Agree - 6 Strongly agree - 7

**Q19. I am in the habit of connecting course material to real-world problems.**

Not Applicable - 0 Strongly disagree - 1 Disagree - 2 More or less disagree - 3 Undecided - 4 More or less agree - 5 Agree - 6 Strongly agree - 7

**Q23. I am in the habit of linking material from different courses and disciplines.**

Not Applicable - 0 Strongly disagree - 1 Disagree - 2 More or less disagree - 3 Undecided - 4 More or less agree - 5 Agree - 6 Strongly agree - 7

**Q25. I am in the habit of linking lab activities with class activities.**

Not Applicable - 0 Strongly disagree - 1 Disagree - 2 More or less disagree - 3 Undecided - 4 More or less agree - 5 Agree - 6 Strongly agree - 7

### Subcomponent 6) Clarity of Instruction: 20, 22

**Q 20. The material is presented in a clear way.**

Not Applicable - 0 Strongly disagree - 1 Disagree - 2 More or less disagree - 3 Undecided - 4 More or less agree - 5 Agree - 6 Strongly agree - 7

**Q 22. I felt comfortable asking questions to the Instructor.**

Not Applicable - 0 Strongly disagree - 1 Disagree - 2 More or less disagree - 3 Undecided - 4 More or less agree - 5 Agree - 6 Strongly agree - 7

